# Temporal dynamics of endogenous attentional modulation without cue awareness

**DOI:** 10.1101/319137

**Authors:** Yuma Osako, Shota Murai, Jun Shimpaku, Kohta I. Kobayasi

## Abstract

Invisible visual stimuli can regulate our broad cognitive performance in the pursuit of current goals. Endogenous spatial attention is an important modulator of cognitive performance, and it can be triggered by unconscious cues. However, how its effect changes with time remains unclear. Here, we show that endogenous attention was triggered by an arrow-cue whose direction participants reported being unaware of but which affected the task performance in a time-dependent manner. Participants were asked to remember the directions of eight Landolt c rings (target memory array) after arrow-cue presentation, which was designed to orient their attention to a certain c ring. Then, we applied a delay, ranging from 83 ms to 1000 ms, between the arrow-cue and the target memory array presentation (the possible delays were equally spaced on a logarithmic scale). The attentional effect was greater for the 83, 183, 250 and 333 ms delays than the other six possible delays. In contrast, its effect was maintained irrespective of the delay when the participants reported being aware of the cue direction. Thus, awareness of arrow-cue direction was necessary to maintain endogenous attentional modulation, and its modulation without arrow-cue direction awareness was limited in a time-dependent manner.

## Introduction

A visual stimulus falling outside our awareness can bias our conscious experience^1,2^ and behavioural responses^3,4^. These phenomena have been reported in blindsight patients who suffer from cortical blindness due to lesions in the primary visual cortex (V1). They can discriminate or respond to visual stimuli even though they cannot consciously see them^5–7^. This unconscious processing of invisible stimuli was also reported in healthy human participants in cases of masking^8^, binocular rivalry^9^, inattentional blindness^10^, motion-induced blindness^11^ and continuous flash suppression^12^. Such unconscious visual information can affect performance in a broad range of cognitive tasks, such as object recognition^13,14^ and decision-making^15–18^. This unconscious processing may accelerate our cognitive and informative life to improve and promote the efficiency of conscious performance.

Spatial attention is one of the main factors that can improve or promote the processing of visual stimuli^19^. Spatial attention is triggered by visual stimuli and is suggested to modulate the neural response at the early and late stages of stimulus processing^20–23^. Unconscious stimuli also trigger the shift of spatial attention^24–28^. However, most studies demonstrated that these effects were elicited by peripheral or spatially compatible cues, which could drive the shift of spatial attention in an exogenous manner. A masked prime presented in a central position can nonetheless affect the shift of spatial attention to the peripheral space^29,30^. These findings indicate that an unconscious prime can be used to handle the shift of spatial attention in an endogenous manner. Clarifying the neural correlates of endogenous spatial attention, which unconscious visual stimuli trigger, is important in understanding the neural mechanisms of unconscious performance. However, little is known about whether the effect of such spatial attention is temporally changed, as well as when the unconscious shift of attention occurs after subliminal visual stimulus presentation. To understand the temporal dynamics of these unconscious modulations in performance, it is necessary to address the temporal dissociation of endogenous attentional effects with and without cue awareness.

Thus, we utilized the effect of spatial attention on encoding visual stimuli into visual short-term memory (VSTM). For example, when participants attend to a specific location in their visual field, the target stimulus at that location is more likely to be encoded into VSTM than other target stimuli at unattended locations^31–33^. Then, to test the temporal dynamics of attentional effects, we applied a pre-cueing paradigm^19^, in which an attentional arrow-cue was presented before a delay of 83-1000 ms until a target memory array presentation (eight Landolt c rings shaping an imaginary circle). The subjects were instructed to encode the visual memory array into VSTM, and following the retention interval (100 ms), a probe cue was presented at the target memory array. The participants answered in which direction the Landolt c ring at that position pointed. The attentional arrow-cue was intended to modify the location of attentional space, and we rendered its arrow direction invisible by using forward and backward masking.

## Methods

### Participants and Apparatus

A total of twenty-two healthy adults (13 males, 9 females, aged 21-27) participated in this study voluntarily. All participants gave informed consent and were paid for their participation. All experiments were performed in accordance with the guidelines for experiments at Doshisha University with the approval of the Research Committee of Doshisha University. All the data were anonymously treated, and the private information was prevented from leakage.

The experiment was programmed in MATLAB using Psychotoolbox^34^ and was conducted in a sound-proof and darkened room with only the monitor producing light. All visual stimuli were generated by a 60 Hz-refresh-rate monitor and displayed at a resolution of 1920 × 1080 (FG2421, EIZO, Ishikawa). The masking stimuli were synthesized with random RGB dots. The masking procedure was one of the common ways to render cue blindness^35,36^. The monitor used a grey background to reduce afterimages. All responses were collected by inputs on a standard keyboard. All participants set their jaw on a chin rest to stabilize their eye position and to keep a viewing distance of approximately 60 cm.

### Stimuli and Procedure

We measured the temporal change in the accuracy in the probe test under two conditions: (1) whether participants reported that they were aware or unaware of the arrow-cue direction and (2) whether the arrow-cue was correctly predictive for later probe position (validity). We presented the arrow-cue at two contrasts (the trial ratio of low to high contrast equalled 4 to 1) to control the masking effect. The low-contrast cue had 81% of the luminance of the high-contrast cue (Fig. 1: Cue). The masking stimulus was a picture of random RGB dots that covered the area where the cue was presented. The target memory array was presented as eight Landolt c rings, and each ring’s slit (ring direction) was positioned in eight different possible directions. The eight directions were in the 12:00, 1:30, 3:00, 4:30, 6:00, 7:30, 9:00 and 10:30 o’clock directions. The ring direction was produced in pseudo-random order from the eight directions. Each ring was located equidistantly at the eight different positions, forming an imaginary circle at <9 ° in diameter from the centre fixation point (Fig. 1: Target memory array). The eight rings locations were the 12:00, 1:30, 3:00, 4:30, 6:00, 7:30, 9:00 and 10:30 o’clock locations. The control cue was a circle filled with the same colour and contrast as the low-contrast cue. The probe cue was a white-bordered square.

**Figure 1.**
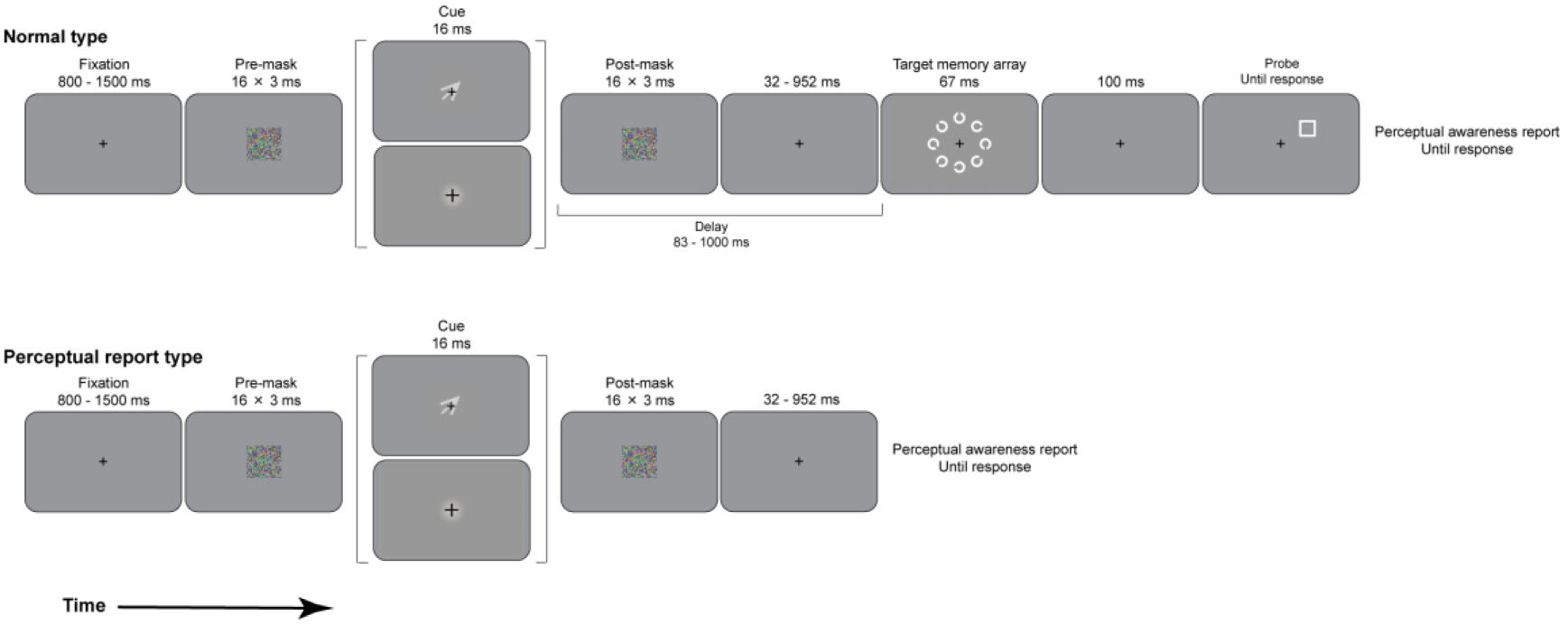
Schematics of sequences of the behavioural paradigm. (Upper) Schematic of a normal trial. In each trial, a fixation cross was presented for 800- 1500 ms, followed by three pre-masks. After the pre-mask presentation for 48 ms, an arrow-cue or circle-cue was presented for 16 ms, followed by post-mask presentation in the same way as the pre-mask presentation. Following a 32-952 ms delay, the target memory array was presented for 67 ms. Participants were instructed to transfer it into their visual short-term memory and answer the probe test after the retention interval (100 ms). After the probe test, they performed an unspeeded perceptual report. (Lower) Schematic of the perceptual report type. The equal sequences before the target memory array presentation in a normal trial. In this trial type, the participants performed only the perceptual report after a 32-952 ms delay.

The experimental paradigm (Fig. 1) adapted forward and backward masking^37^ based on the experimental 1 diagram of Delvenne and Holt (2012). A similar method has been used in several previous studies^29,33^. Our paradigm had two trial types: normal type and perceptual report type. In the former, participants performed probe tests and reported their perceptual awareness, whereas in the latter, they only reported their perceptual awareness to ensure its quality in the normal trial type.

All participants had some practice sessions and six test sessions. In the practice sessions, the participants trained to understand the behavioural paradigm. To help them understand the experimental paradigm, we used only high-contrast and valid arrow-cues in practice sessions. In test sessions, low contrast arrow-cues were used with 80% probability, and the remaining cues were high contrast, with a mixture of valid, invalid and control arrow-cues (ratio of valid:invalid:control trials = 3:1:1). Each session included 100 trials and usually took 1.5 hours for the six test sessions for one participant.

In normal trials, each trial started with a central fixation cross that was presented for 800–1500 ms. Following the fixation cross, three pre-masks were presented for 48 ms (each 16 ms). After the pre-mask presentation, an arrow-cue directed toward one of eight locations was presented for 16 ms, followed by three post-masks (same as pre-mask presentation). The target memory array was then presented for 67 ms after a 32–952 ms delay following the flashing of the post-masks, followed by a 100 ms blank interval and by a probe that remained present until the response key was pressed. After the probe response, participants made an unspeeded response to report their perceptual awareness of the arrow-cue, whether the direction of the arrow-cue was seen clearly and was discriminable (fully visible), seen obscurely and was indiscriminable (fragmented visible), or not seen at all (invisible), by pressing the G, H or J key, respectively.

The perceptual awareness report trial had the same procedure before the target memory array presentation. After the delay, participants responded with their unspeeded perceptual awareness report without the probe test.

The conditions were randomly changed during the experiment. All trials started after pressing the keyboard in the perceptual awareness report of the previous trial. In one session (100 trials), a normal trial type included 90 trials, with 54 trials counted as valid, 18 trials as invalid, 18 trials as control arrow-cue presentations, and the remaining 10 trials as perceptual awareness trials. The delay was selected randomly and independently with a duration of 83, 116, 150, 183, 250, 333, 433, 566, 750, or 1000 ms (equally spaced on a logarithmic scale).

## Results

First, the accuracy in the probe test irrespective of the delay was calculated (Fig. 2A). We performed Tukey’s pair-wise post hoc test after two-way analysis of variance (ANOVA) with awareness of arrow-cue direction and three arrow-cue direction types (validity and control). The participants performed significantly better in valid-aware and valid-unaware conditions compared with control and invalid conditions (p<0.001 and p<0.001, respectively). In contrast, accuracy did not differ significantly between the control and two invalid conditions. The reaction time was only significantly shorter in the valid-aware condition compared to the other conditions (Fig. 2B, p = 0.004 with valid-unaware, p = 0.001 with control and invalid-aware and p < 0.001 with invalid-unaware).

**Figure 2.**
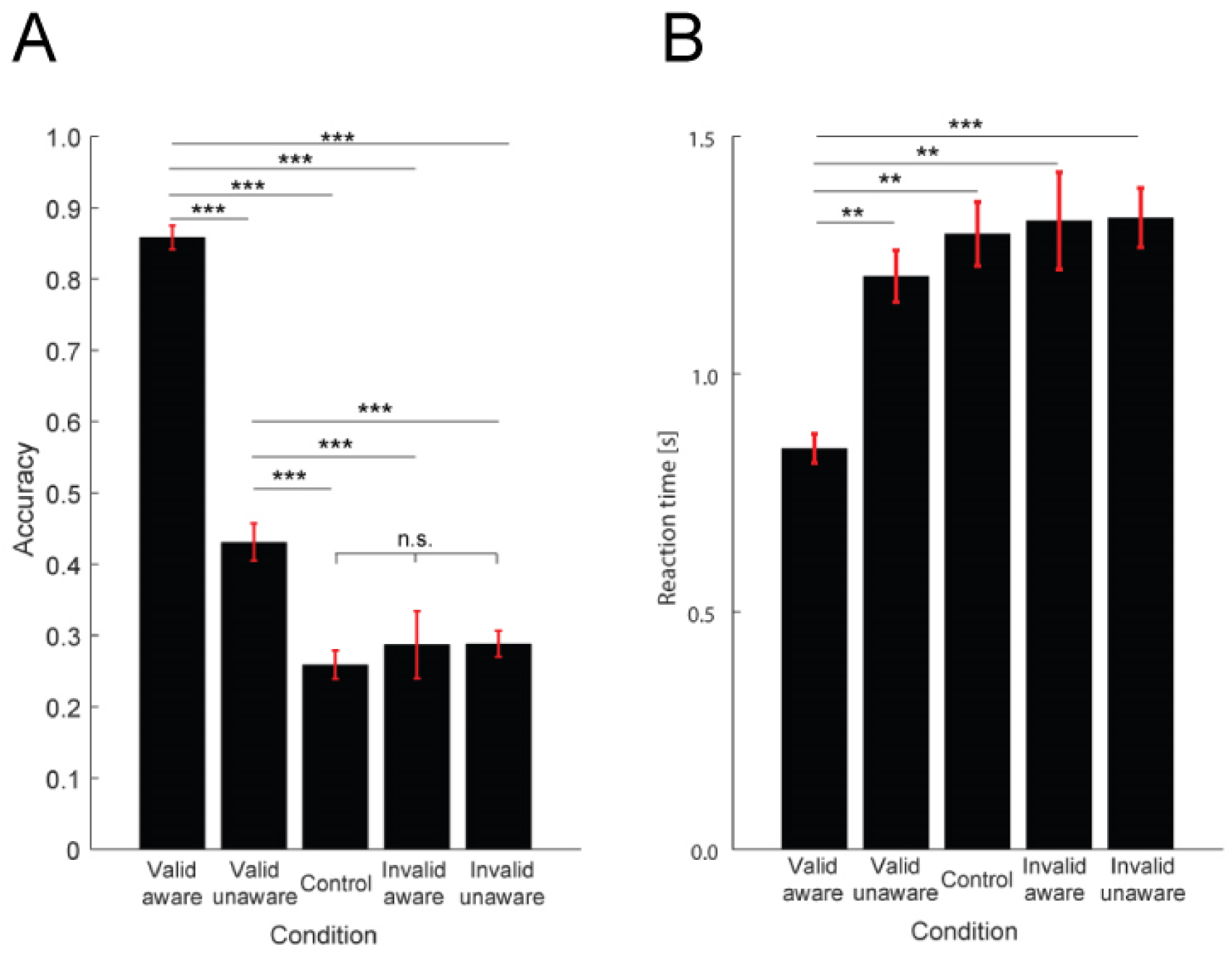
Behavioural performance of the pre-cue paradigm. (A) The accuracy of the probe test is shown as the mean ± standard error of the mean (SEM). (B) The reaction time of the probe test is shown as the mean ± SEM. ^**^p<0.01, ^***^p<0.001.

We therefore explored the relationships of the accuracy and reaction time with the delay, i.e., the inter-stimulus interval between the presentation of the arrow-cue and target memory array (Fig. 3A-B). The accuracy was 75–90 % in the valid-aware condition (Fig. 3A, blue line) and gradually increased upon increasing the delay in the control condition (Fig. 3A, black line). In contrast, time-dependent changes were observed in the other conditions (Fig. 3A, red, purple and green lines). The reaction time in the valid-aware condition was significantly shorter regardless of the delay compared to the other conditions, but the other conditions did not differ significantly based on the delay (Fig. 3B).

**Figure 3.**
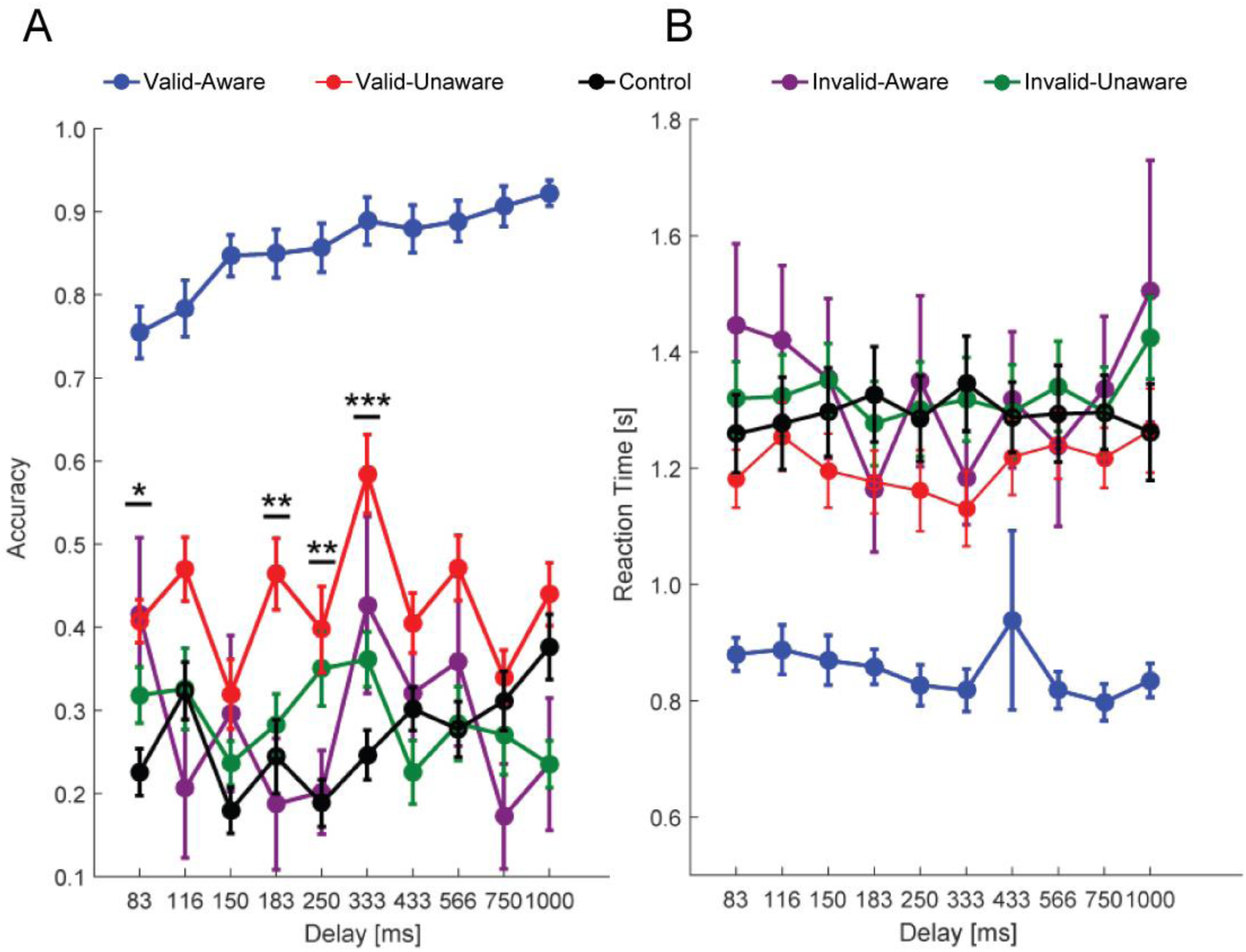
Temporal changes in behavioural performance. (A) Temporal change in accuracy in the probe test. Each line shows the mean ± SEM. (B) Temporal change in reaction time in the probe test. Each line shows the mean ± SEM. ^*^p<0.05, ^**^p<0.01, ^***^p<0.001.

We next addressed the temporal dynamics of the endogenous attentional effect on behavioural performance. We defined delta as the difference in accuracy between the valid-unaware and control (the value of valid-unaware minus control). The accuracy exhibited a clear peak at approximately 333 ms after presentation of the arrow-cue (Fig. 4A). The reaction time also had the lowest delta for the 333 ms delay (Fig. 4B). We then analysed the correlation between these deltas, and there was a significant negative correlation (Fig. 4C, Pearson’s r = -0.768, p = 0.010).

Finally, we classified the delay time into three phases to analyse the time-dependent changes in longer time windows: 83-150, 183-333 and 433-1000 ms. The classification sizes were based on the significantly better accuracy between valid-aware and control conditions in three consecutive delay times (183, 250 and 333 ms, see Fig. 3A). Accuracy was significantly greater in the intermediate phase compared to the first and last phases (Fig. 4C, p = 0.024 vs. first phase and p < 0.001 vs. last phase). The reaction time in the intermediate phase was significantly decreased compared to the last phase (Fig. 4D, p = 0.014) and was also decreased compared to the last phase with a significant trend (Fig. 4D, p = 0.069).

**Figure 4.**
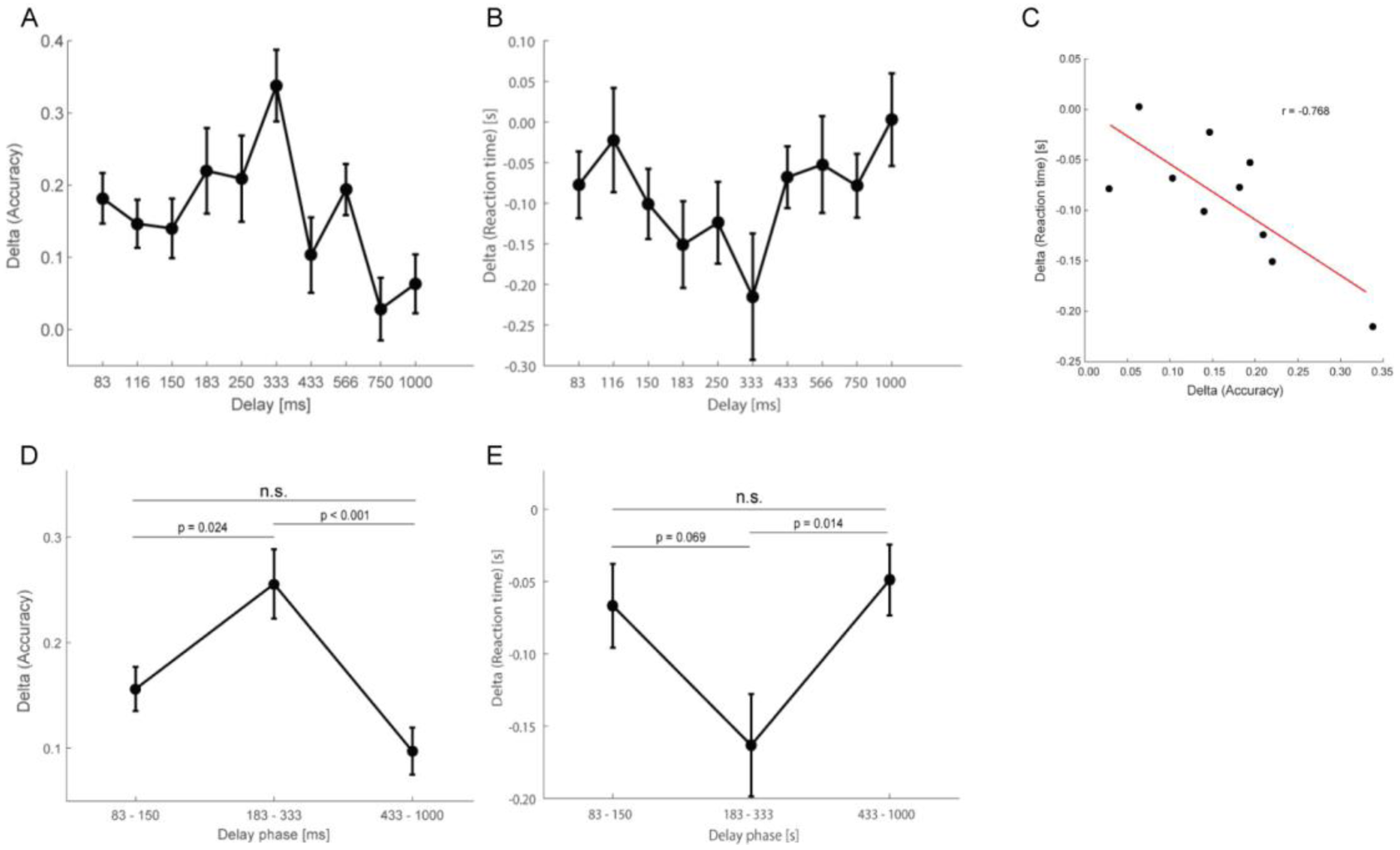
Temporal changes in the delta. (A) Temporal change in the delta of the accuracy is shown as the mean ± SEM. (B) Temporal change in the delta of the reaction time is shown as the mean ± SEM. (C) Significant correlation between the delta of the accuracy and the delta of the reaction time (Pearson’s correlation, r=-0.768, p=0.010). (D) Temporal change in the delta of the accuracy in large time windows is shown as the mean ± SEM. (E) Temporal change in the delta of the reaction time in large time windows is shown as the mean ± SEM.

## Discussion

Many previous studies exploring the unconscious modulation of our cognitive performance by unconscious visual information have focused on the dissociation between reports of subjective cue awareness and actual performance. The quest for neural mechanisms for the unconscious modulation implies that how the dissociation changes through the specific time windows is quite important to understand the neural representation or processing of it.

As an approach to answer this question, we examined the temporal dynamics of the endogenous attentional effect by measuring the transfer of the visual stimulus to VSTM while manipulating the awareness of arrow-cue directions. Our results reveal that endogenous attention was elicited irrespective of cue awareness, as previous studies reported (Fig. 2A), and significantly better performance was observed with 83, 183, 250 and 333 ms delays when participants reported being unaware of arrow-cue direction (Fig. 3A). We also analysed the dissociation distance (i.e., delta) between valid-unaware and control conditions using accuracy and reaction time as measures of the endogenous attentional effect. The delta of the accuracy formed the shape of an arch, with a peak at 333 ms (Fig. 4A). The delta of the reaction time showed the reverse trend, with the lowest value at 333 ms (Fig. 4B). There was a significant negative correlation between these deltas (Fig. 4C, Pearson’s r = -0.768, p = 0.010). This indicates that the endogenous attention affects cognitive performance in a time-dependent manner.

To address its time-dependent manner, we divided the delays into three phases due to the significantly better accuracy in the 183, 250, and 333 ms delay conditions (Fig. 3A). The intermediate phase showed significant differences compared to first and last phases (Fig. 4D, E). This suggests that endogenous attention elicited by an unconscious arrow-cue could affect performance during specific time windows. In contrast, endogenous attention elicited by a conscious arrow-cue maintained its effect regardless of the delay. These results suggest that endogenous attentional modulation was stronger at 183-333 ms delays and did not occurred at longer delays (433 ms and longer) in this study.

One model for explaining the different time-dependent results between valid-aware, valid-unaware and control conditions is a cross of temporal expectation and attentional scope. Temporal expectation has often been considered a hazard function of the presentation of the imminent target stimuli^38,39^. In our study, a gradual increase in the accuracy with increasing delay was observed in valid-aware and control conditions. This implies that participants could manipulate their attention to target memory array presentation with expectancy because the later delay caused a higher incidence of target presentation in their expectation. Visual attentional scope for the visual local field enhances the visual processing^40,41^. The scope was only covered with a Landolt c ring instructed by the arrow-cue in the valid-aware condition, while the scope was covered with the target memory array in the control condition. This difference in visual attentional scope could grade the accuracy in the valid-aware condition. Then, the temporal dynamics of accuracy in the valid-unaware condition could be expected to be located intermediately between the valid-aware and control conditions, with a gradual increase. However, a gradual increase in accuracy was not observed in the valid-unaware trials in our study. This suggests that attentional modulation, triggered by unconscious arrow-cue direction, has different functional attributions from the cross of temporal expectancy and visual attentional scope.

Finally, the present study supports the notion that endogenous attention is occurred by unconscious visual stimuli^29,30^. Functional brain imaging studies have shown that endogenous visual attention modulates visual processing^42,43^. Our findings and these previous studies offer the opportunity to explore the temporal process of the endogenous visual attention with consciousness of the visual cue-stimulus as an independent variable. Such experiments may shed light on the neural correlates of consciousness^44^.

## Acknowledgements

This research was supported by Research Grant KAKENHI 17H01769 (to K.K.) and 15J04452 (to S.M.). We thank Edward William Ko Uy for helpful comments on the manuscript. We would like also to thank Yoshio Sakurai for helpful discussion.

## Author contributions

Y.O., S.M., J.S. and K.K. designed the experiments. Y.O., S.M. and J.S. performed the experiments and analysed the data. K.K. supervised the project. All authors contributed to discussion. Y.O. and K.K. wrote the manuscript.

